## Supplementary material for "Fast and Flexible Sequence Induction In Spiking Neural Networks Via Rapid Excitability Changes" (from another version of this preprint: version 1, 2018-12-13)

### SUPPLEMENTARY FIGURES

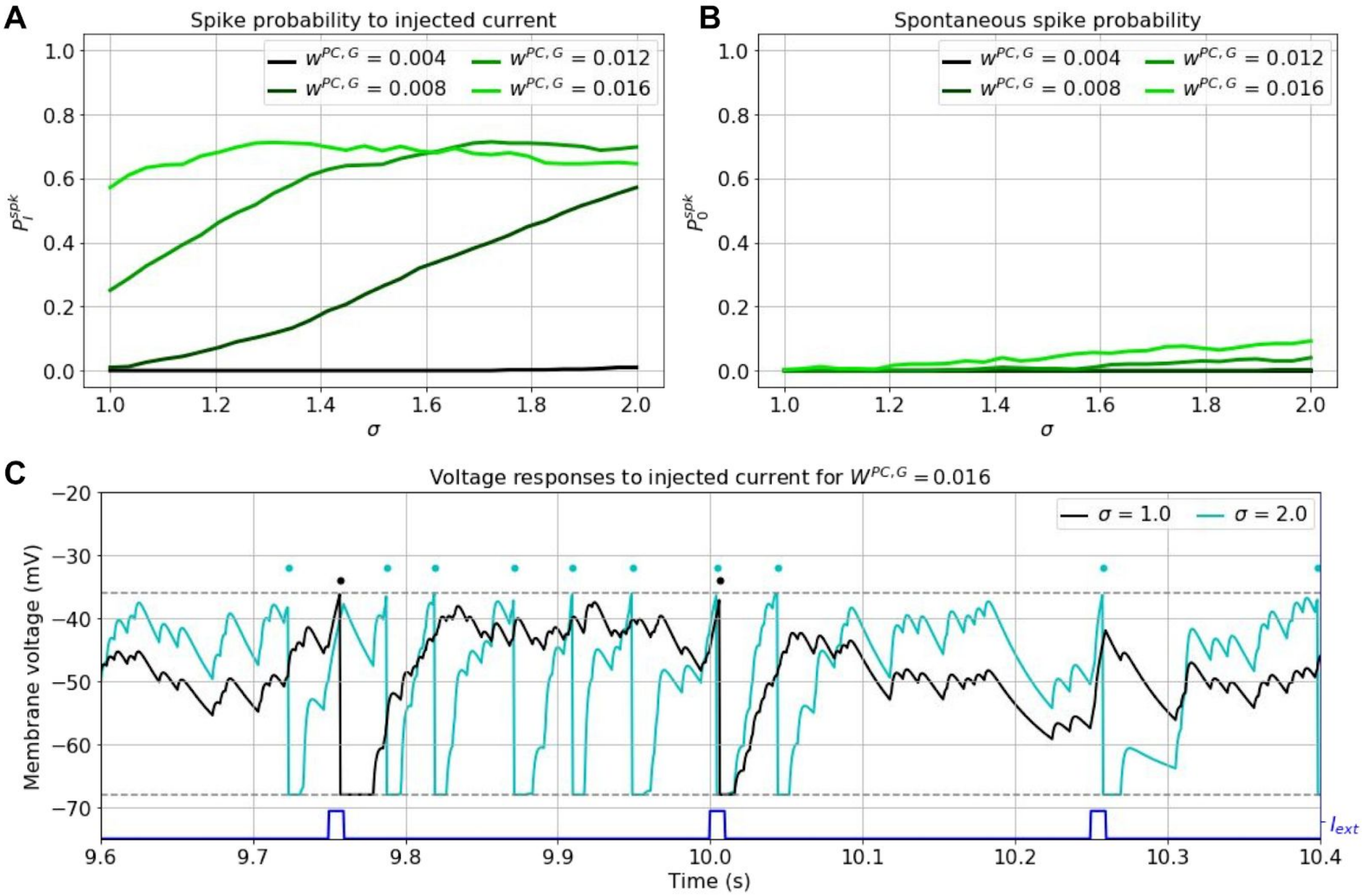


***Fig 1S1: Effects of current pulses on spiking under LTP-IE**. A. Probability of current-evoked spiking for different $w^{PC,G}$ and σ. B. Spontaneous spiking probability for different different $w^{PC,G}$ and σ. C. Example voltage traces for neurons with two different σ given large $w^{PC,G}$. Same format as **Fig 1F**.*

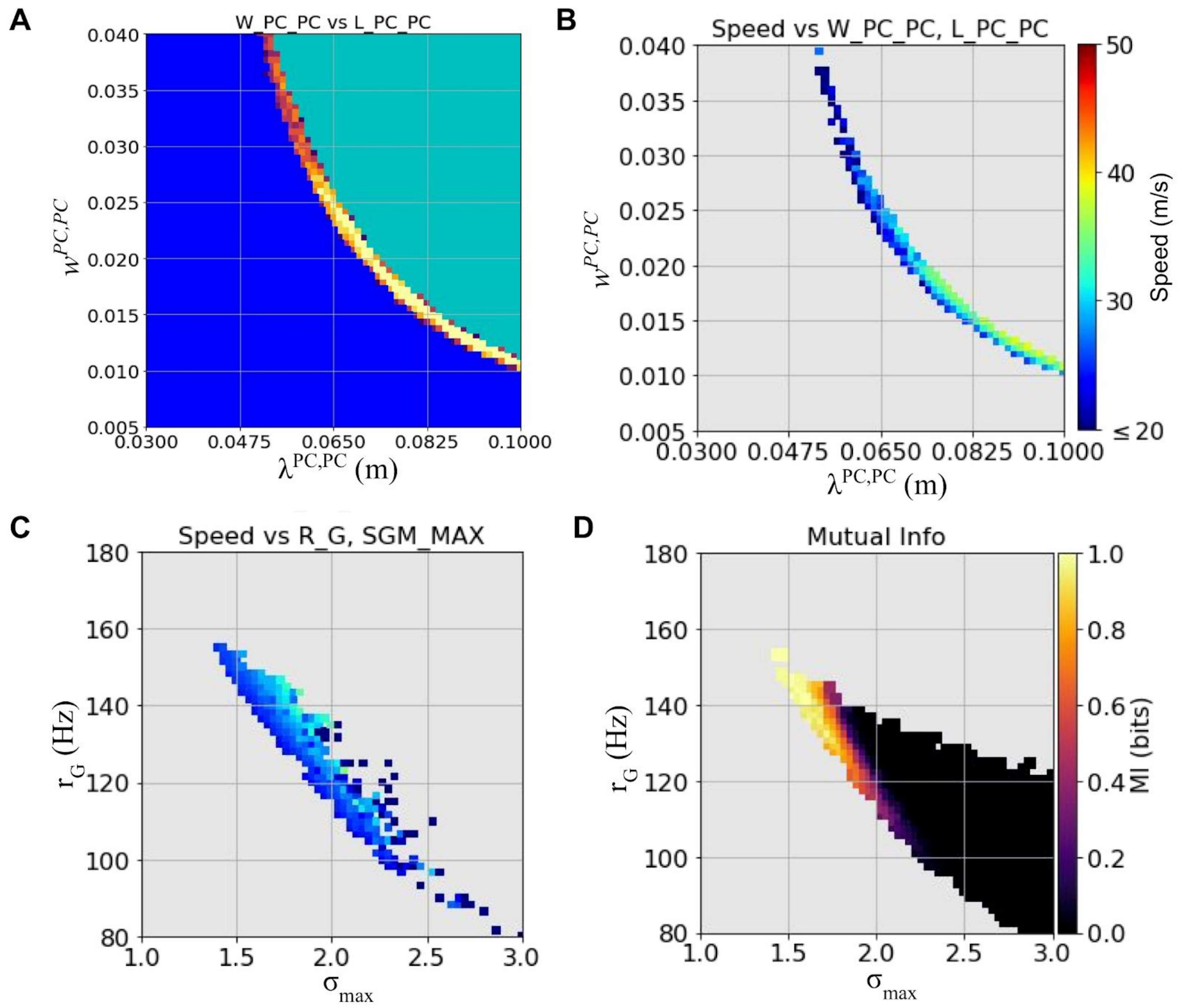


***Fig 3S1: Replay features for denser networks****. A. As in* ***Fig 3D*** *but for a "dense" network of 2000 neurons. B. As in* ***Fig 3J*** *but for dense network. C. As in* ***Fig 3K*** *but for dense network. D. As in* ***Fig 3L*** *but for dense network.*

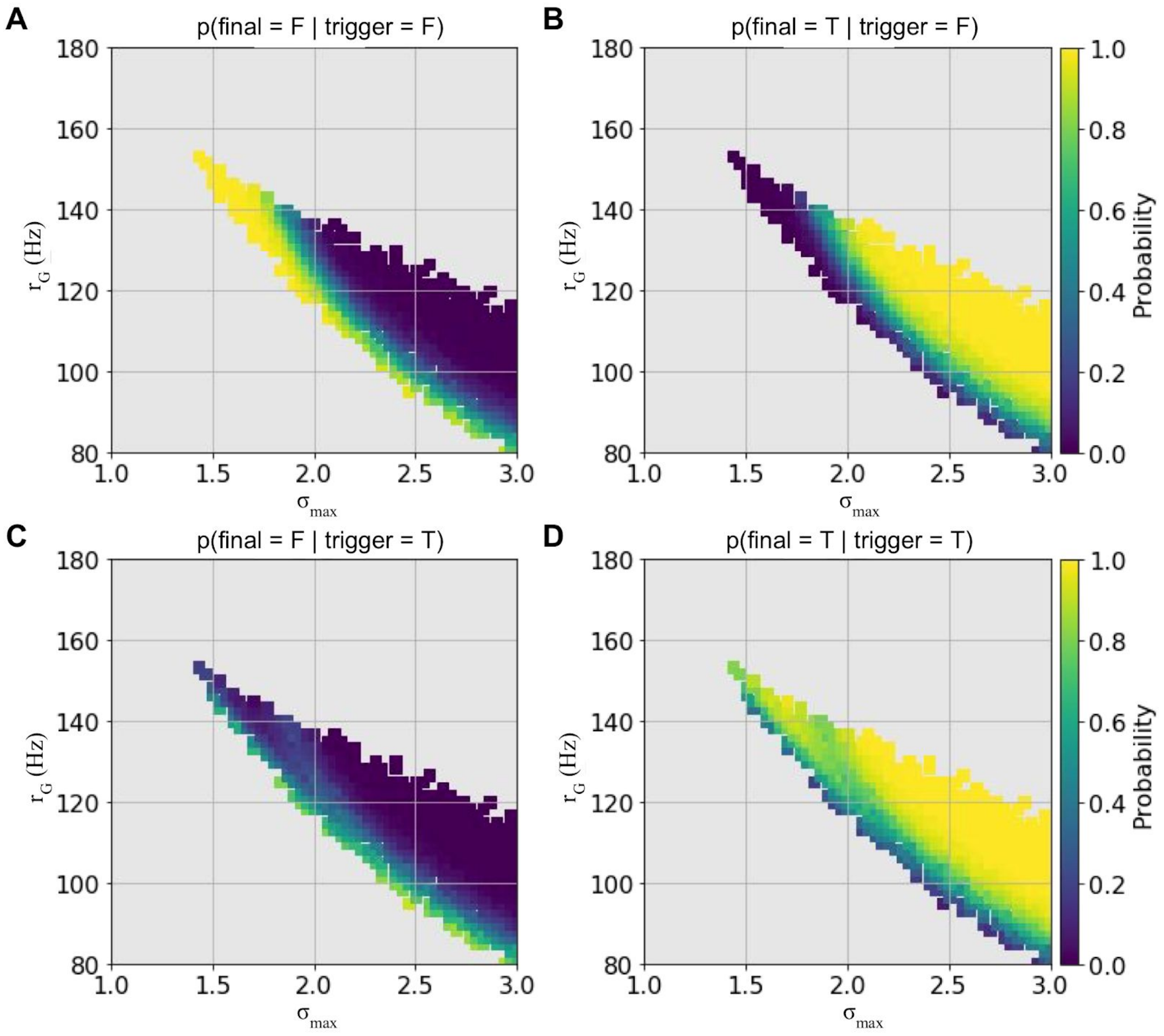


***Fig 3S2: Mutual information calculation**. Plots of conditional probabilities used to calculate mutual information in **Fig 3L**. A. Probability of no elevated spiking at opposing end of trajectory ("final = F") given no initiating trigger ("trigger = F"). B. Probability of elevated spiking at opposing end of trajectory ("final = T") given no initiating trigger. C. Probability of no elevated spiking at opposing end of trajectory given initiating trigger ("trigger = T"). D. Probability of elevated spiking at opposing end of trajectory given initiating trigger.*
